# Transition of Wnt signaling microenvironment delineates the squamo-columnar junction and emergence of squamous metaplasia of the cervix

**DOI:** 10.1101/443770

**Authors:** Cindrilla Chumduri, Rajendra Kumar Gurumurthy, Hilmar Berger, Stefanie Koster, Volker Brinkmann, Uwe Klemm, Hans-Joachim Mollenkopf, Hermann Herbst, Mandy Mangler, Thomas F Meyer

**Affiliations:** Department of Molecular Biology, Max Planck Institute for Infection Biology, Berlin, Germany; Institute of Pathology, Vivantes Klinikum Berlin, Rudower Straße 48, 12351 Berlin, Germany; Charité University Medicine, Department of Gynecology, Berlin, Germany; Auguste-Viktoria-Klinikum, Vivantes, Klinik für Gynäkologie und Geburtsmedizin 12157 Berlin, Germany

## Abstract

The transition zones (TZ) between squamous and columnar epithelium constitute hotspots for the emergence of cancers. Carcinogenesis at these sites is often preceded by the development of metaplasia, where one epithelial type invades the neighboring one. It remains unclear how these niches are restrained at the boundary between the two epithelial types and what factors contribute to metaplasia. Here we show that the cervical squamo-columnar junction derives from two distinct stem cell lineages that meet at the TZ. In contrast to the prevailing notion, our analysis of cervical tissue showed that the TZ is devoid of any locally restricted, specialized stem cell population, which has been implicated as precursor of both cervical squamous cell carcinoma and adenocarcinoma. Instead, we reveal that these cancers originate from two separate stem cell lineages. We show that the switch in the underlying Wnt signaling milieu of the stroma is a key determinant of proliferation or quiescence of epithelial stem cell lineages at the TZ. Strikingly, while the columnar lineage of the endocervix is driven by Wnt signaling, the maintenance of squamous stratified epithelium of the ectocervix and emergence of squamous metaplasia requires inhibition of Wnt signaling via expression of Dickkopf2 (Dkk2) in the underlying stroma. Moreover, Notch signaling is required for squamous cell stratification. Thus, our results indicate that homeostasis at the TZ is not maintained by a transition from one epithelial type to another but rather results from alternative signals from the stromal compartment driving the differential proliferation of the respective cell lineages at the squamo-columnar junction.

## Introduction

Recent years have seen rapid progress in our understanding of how adult epithelial tissues are maintained by dedicated stem cell niches. In many cases, Wnt signaling initiated from the underlying stroma plays decisive roles. What is entirely unclear so far is how these niches are restrained at the boundary between two different types of epithelia, e.g. columnar and stratified ones. Such transition zones are found in several organs in the human body, e.g. the gastro-esophageal junction, as well as the cervix. These sites almost invariably show a predisposition towards transformation, which is preceded by metaplasia, where one epithelium type invades another.

Cervical cancer predominantly occurs at the TZ, where the stratified squamous epithelium of the ectocervix and the simple columnar epithelium of the endocervix meet ^1,2^. It is one of the most common and deadly cancers in women and occurs as two histologically distinct types: adenocarcinomas (ADC) and squamous cell carcinomas (SCC) ^3^. Both ADC and SCC are proposed to originate from a common precursor stem cell population with a unique immunophenotype that is localized exclusively at the TZ. This precursor cell population is proposed to consist either of residual embryonic cytokeratin 7 (KRT7) positive cells ^4^, p63+/KRT7+/KRT5+ transitional basal cells ^5^, or reserve cells ^6^. As a result, prophylactic ablation of residual embryonic cytokeratin 7 (KRT7) positive cells has been proposed as a method to prevent cervical cancers ^7,8^. However, how this population gives rise to two epithelial types and the molecular mechanisms that govern maintenance of the TZ and define the distinct epithelial compartments has not been studied. In addition, the changes that contribute to metaplasia at the TZ of the cervix (or indeed other tissues) also remain unclear.

Here, using *in vivo* lineage tracing, cervical organoid models, single molecule RNA in situ hybridization (RNA-ISH) and a mouse model of squamous metaplasia, we show that endo-and ectocervical epithelia are derived from two separate lineage-specific stem cell populations that meet at the TZ. Using bioinformatic analysis as well as immunohistochemistry of cancer tissue, we find that the transcriptional signatures of these lineages correspond to those of SCC and ADC, indicating that these two histologically distinct cancer types also arise from the two different lineages rather than from a common precursor. We show that two distinct active or inhibitory Wnt signaling milieus are established in the endo-and ectocervical region by Wnt-pathway agonists and antagonists, respectively, which are differentially expressed in the epithelium and sub-epithelial stroma on either side of the TZ *in vivo*. Organoids derived from these two distinct stem cell lineages show a strong divergence in their requirement for Wnt signaling. Strikingly, we demonstrate that endocervix also harbors squamous stem cells, which are kept in a quiescent state by the Wnt signaling microenvironment that promotes proliferation of the columnar lineage. *In vitro*, these quiescent stem cells can give rise to squamous epithelium in the absence of Wnt. This is the first time that the presence of two stem cell lineages with opposing signaling requirements within the same niche has been observed. In addition, using an *in vivo* mouse model of cervical metaplasia, we show that loss of Wnt signals by induction of Wnt inhibitory signaling molecules in the endocervical region leads to the outgrowth of the pre-existing ectocervical stem cells within the transition zone and the endocervix. These findings provide the first mechanistic underpinning of how homeostasis is maintained at the transition zone and how development of metaplasia is initiated.

## Results

### Distinct cellular origins of squamous and columnar epithelium

To obtain deeper insight into the cellular composition and molecular determinants of TZ maintenance, we carried out a detailed analysis of marker profiles in human and mouse cervical epithelium. Strikingly, our comprehensive, unbiased analysis including the entire endo-and ectocervix regions failed to detect any specific cell type that was restricted exclusively to the TZ - in contrast to the prevailing concept. Instead, we observed two distinct epithelial lineages, with KRT5 expressed throughout the squamous stratified epithelium and KRT7/KRT8 expressed throughout the columnar epithelium. At the TZ there is an overlap of both lineages, where KRT5+ basal cells appear to displace overlying KRT7+/KRT8+ columnar cells to form squamous stratified epithelium (Fig. 1 A-D and Fig. S1 A-D). RNA-ISH confirmed that KRT8 and KRT5 expression was restricted to the columnar epithelium and basal/parabasal cells of squamous epithelium, respectively (Fig. 1 E-F, and Fig. S2 A-B). In contrast to previous reports describing a discrete KRT7 population, restricted to the TZ ^4,5^, we observed KRT7 expression at high levels throughout the endocervical epithelium and sparse expression in the ectocervical epithelium (Fig. 1 G and Fig. S2 C). We also observed that patches of subcolumnar KRT5+ cells occurred sporadically beneath KRT7+/KRT8+ cells within the endocervix (Fig. S1A). These islands of KRT5+ cells may correlate with foci of squamous metaplasia that are frequently observed within the endocervix and account for 10% of premalignant squamous intraepithelial lesions (SIL) ^9,10^.

**Figure 1.**
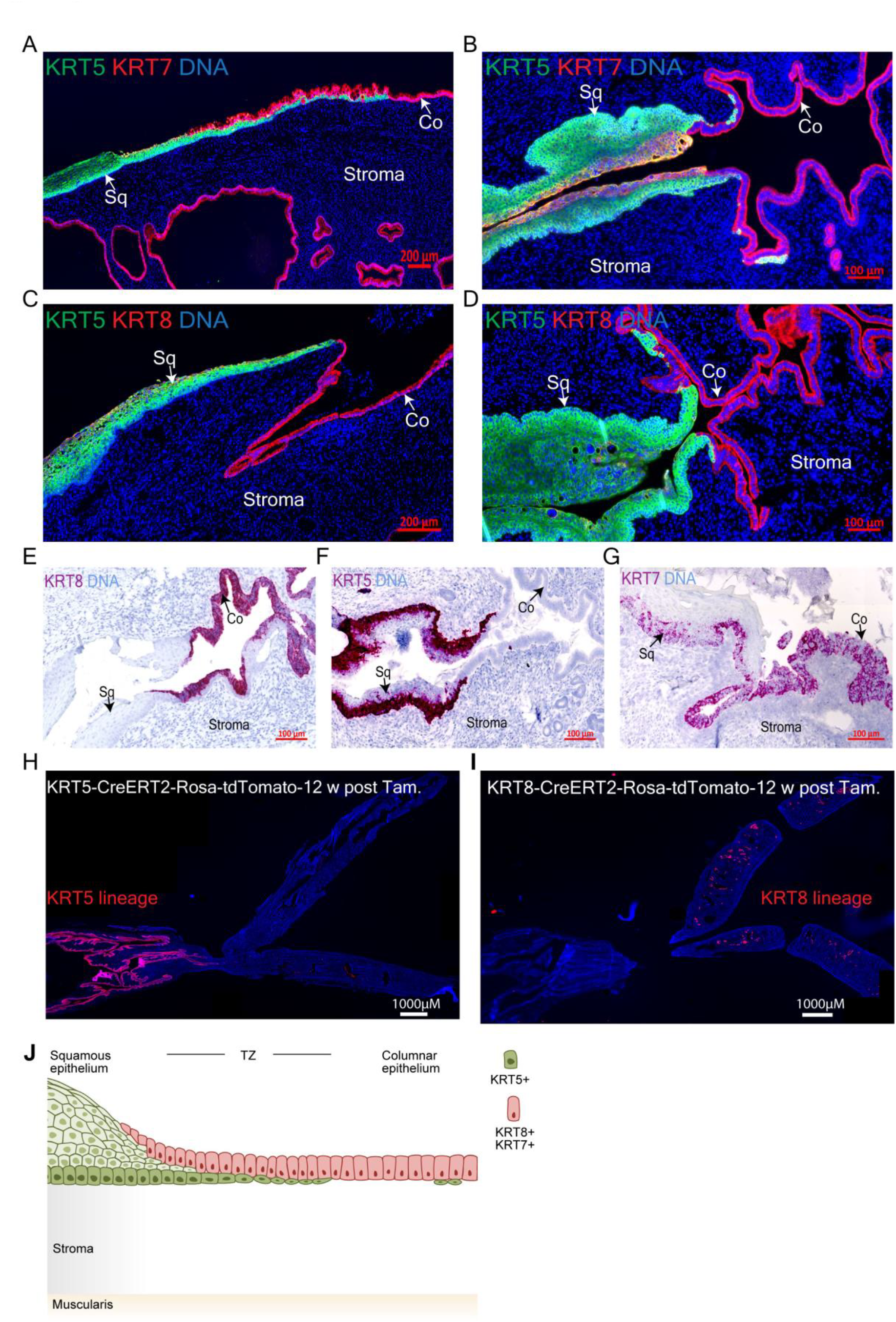
Cervix consists of two distinct KRT5+ stratified and KRT7+/8+ columnar epithelial lineages. Transition zone (TZ) including stratified and columnar epithelium from human (A,C) and mouse (B, D) cervix tissue sections immunolabeled with antibodies against KRT5, KRT7, and KRT8; nuclei are shown in blue. (E-G) Single molecule RNA ISH brightfield images of mouse cervix TZ for KRT8, KRT5 and KRT7; nuclei are shown in blue. (H-I) Tiled images of tissue sections from the genital system of 16 week-old KRT5CreErt2/Rosa26-tdTomato and KRT8CreErt2/Rosa26-tdTomato mice after tamoxifen induction at the age of 4 weeks. (J) Schematic of the stratified and columnar lineages and the TZ of the cervix. Tiled images were acquired with AxioScan imager and are representative of *n* = 3 biological replicates. Arrows indicate squamous epithelium (Sq) and columnar epithelium (Co).

Further, using KRT5CreErt2/Rosa26-tdTomato and KRT8CreErt2/Rosa26-tdTomato mice for genetic lineage tracing we confirmed the presence of two distinct epithelial lineages in the cervix (Fig. 1 H-I and J). In these mice, Cre was induced by tamoxifen injection 4 weeks after birth. At 16 weeks of age, KRT5+ cells exclusively populated the stratified epithelium, including all differentiated cells, while KRT8+ cells exclusively generated endocervical epithelium.

### Wnt agonists and antagonists in epithelia and stroma orchestrate TZ

To gain insight into the factors that regulate the two lineages and maintain the TZ, we established and defined conditions that facilitate long-term *in vitro* propagation of ecto-and endocervical epithelial stem cells as 3D organoids. Wnt signaling was described to be essential for generation and long-term maintenance of adult epithelial stem cell-derived organoids from the various tissues so far described, as shown by the requirement of Wnt3a and R-spondin 1 in the tissue-specific organoid culture media ^11^. In contrast to this, the presence of Wnt3a and R-spondin 1 (RSPO1) in the culture medium was detrimental for the formation and expansion of human and mouse squamous stratified organoids derived from single ectocervical stem cells (Fig. 2A and Fig. S3A-C). The presence of epidermal growth factor (EGF), fibroblast growth factor 10 (FGF-10), and the inhibition of transforming growth factor beta (TGF-β) and bone morphogenetic protein (BMP) signaling, on the other hand, was essential for long-term maintenance of these organoids. Squamous stratified organoid growth was further increased in the presence of the cAMP pathway agonist forskolin (FSK) (Fig. 2A and Fig. S3D-E). These organoids are KRT5+/KRT7-and fully recapitulate the *in vivo* tissue architecture, with stratified epithelial layers decorated with the adhesion molecule E-cadherin (CDH1) (Fig. S3F and H). The outer layer consists of p63+ basal cells, which differentiate into parabasal cells with p63 staining fading out towards the lumen. Also, typical of ectocervical tissue, only basal cells express the proliferation marker Ki67 (Fig. S3G).

**Figure 2.**
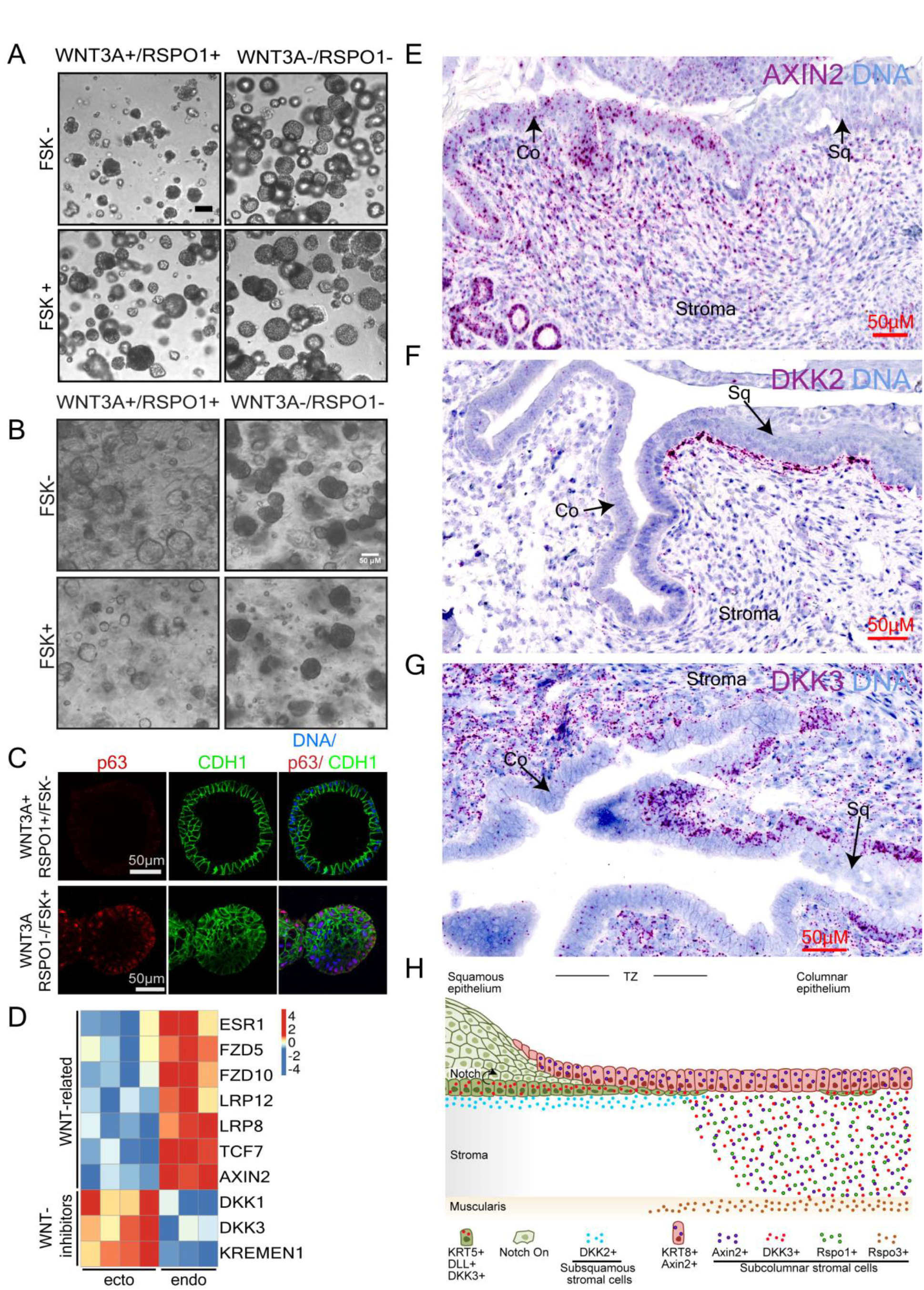
Wnt signaling pathway agonists and antagonists play a key role in ecto and enodcervical development. (**A**) Bright-field images of human ectocervical organoids. Efficient organoid formation depends on absence of Wnt3a and RSPO1 and presence of FSK. (**B**) Cells isolated from endocervical tissue grown in Matrigel with different factors. Wnt signaling is essential for columnar organoid formation, while absence of Wnt drives formation of squamous stratified organoids. (**C**) Columnar and stratified organoids derived from endocervix, containing p63^-^ (columnar) and p63^+^ (stratified) cells. Both express the epithelial marker E-cadherin (CDH1). *n* = 5 biological replicates. (**D**) Expression analysis of differentially regulated genes in human ecto-vs endocervical organoids. Wnt-related genes are expressed at higher levels in endocervical, Wnt inhibitors at higher levels in ectocervical organoids. Columns = biological replicates. (**E-G**) Single molecule RNA ISH of mouse TZ for (**E**) AXIN2, (**F**) DKK2, (**G**) DKK3, nuclei are shown in blue. Tiled images were acquired with AxioScan imager and are representative of *n* = 3 biological replicates. (H) A schematic representation of the distinct epithelial lineages and the underlying tissue microenvironment at the TZ. Arrows indicate squamous (Sq) and columnar (Co) epithelium.

In contrast to the ectocervix, stem cells derived from both proximal and distal endocervix give rise to organoids consisting of a simple columnar epithelial layer when cultured in the presence of Wnt proficient medium containing Wnt3a and RSPO1 (Fig. 2B-C). These organoids faithfully resemble the *in vivo* endocervical epithelium, are KRT7+/KRT5-and exhibit sporadic Ki67 staining (Fig. S4A and S3I). Their self-renewal capacity in culture can be maintained for more than seven months (Fig. S4B). Further, transcriptional profiling of organoids derived from human ecto-and endocervix revealed distinct keratin expression patterns (Fig. S3J).

Strikingly, if cells derived from endocervix tissue were cultured in Wnt deficient medium (FSK+ medium without Wnt3a or RSPO1), they gave rise to p63+ stratified organoids, resembling those derived from ectocervix (Fig. 2B-C, S4C). Since the formation of columnar rather than squamous organoids from endocervical stem cells was dependent on supplementation with Wnt agonists, we investigated the source of Wnt signaling in the cervix. Microarray analysis of organoids and RNA-ISH showed that the transcriptional regulation of Wnt in the endocervix diverges from that in the ectocervix: Wnt agonists are upregulated in columnar epithelium, while the Wnt antagonists Dickkopf WNT signaling pathway inhibitor 3 (DKK3), DKK1 and KREMEN1 are upregulated in squamous epithelium (Fig. 2D, E, G).

Further, we observed that the spatial distribution of extrinsic Wnt agonists and antagonists in the underlying stroma defines the borders between the two epithelial types. Both the Wnt agonist RSPO1 and its downstream target Axin2 are highly expressed in the lamina propria (stroma) beneath the columnar epithelium and RSPO3 in the muscularis of the endocervix (Fig. 2E, S4D-F). Notably, the Wnt antagonist DKK2 is specifically expressed in stroma proximal to the basal cells of the ectocervical squamous epithelium, which express high levels of DKK3 (Fig. 2F-H, S4G-H). In contrast to the ectocervix, high levels of DKK3 expression were observed in the endocervical stroma, while expression of DKK1 was negligible in either region of the cervix (Fig. S5A). Expression levels of DKK4, RSPO2 and RSPO4 also did not show notable regional variation (Fig. S5B-D). Thus, the epithelium of the cervix is maintained by two distinct stem cell populations whose fate is determined by opposing Wnt signaling microenvironments, which are established through the interplay of the epithelial and stromal compartments of the endo-and ectocervix respectively, with a defined switch at the TZ.

### Wnt antagonists, Notch and EGFR signaling maintain ectocervical stemness and differentiation

Next, we sought to identify the cellular pathways that control self-renewal and differentiation in human ectocervical tissue. Microarray analysis showed that squamous ectocervical organoids have a higher expression of Notch-related genes than organoids derived from the endocervical columnar epithelium (Fig 3A). We thus carried out a comparative analysis of 2D cells (2D-ecto), three-day-old early organoids (EO-ecto), and two-week-old, mature differentiated organoids (DO-ecto). 2D cultures were enriched for CDH1+ and p63+ cells, with >60% and >30% of cells showing organoid-forming potential at passage 1 and 8, respectively (Fig. 3B, Fig. S6A). Early organoids consist of 8-16 cells that are undifferentiated and positive for Ki67 and p63 (Fig. 3C and Fig. S6B). Mature organoids consist of several stratified differentiated layers with more than two-thirds of cells differentiated and less than one-third of cells proliferating (Fig. 3D and Fig. S6B). Gene expression patterns of cells from 2D cultures and early organoids show high similarity and display a distinct set of differentially expressed genes compared to mature organoids (Fig. 3E, Table S1-2). Recent studies reported that stem cells from diverse tissue types show similar transcriptional signatures compared to the large divergence observed in the ensuing differentiated tissues ^12^. Comparative analysis of the ectocervical cells (either 2D or EO) and differentiated cell expression profiles to that of frequently upregulated genes in stem cells from diverse tissue types confirmed a high similarity shared with the ectocervical 2D cells and EO in contrast to DO-ecto cells (Fig. 3F, Table S1). This is further supported by the expression profile of genes that are concordantly up-or downregulated in ectocervical 2D-ecto and EO-ecto vs. DO-ecto cells with those of ground state stem cells derived from different tissue types vs. their respective differentiated cells (Fig. S6C, Table S1). Thus 2D-ecto and EO-ecto define characteristics of ectocervical stem cells.

**Figure 3.**
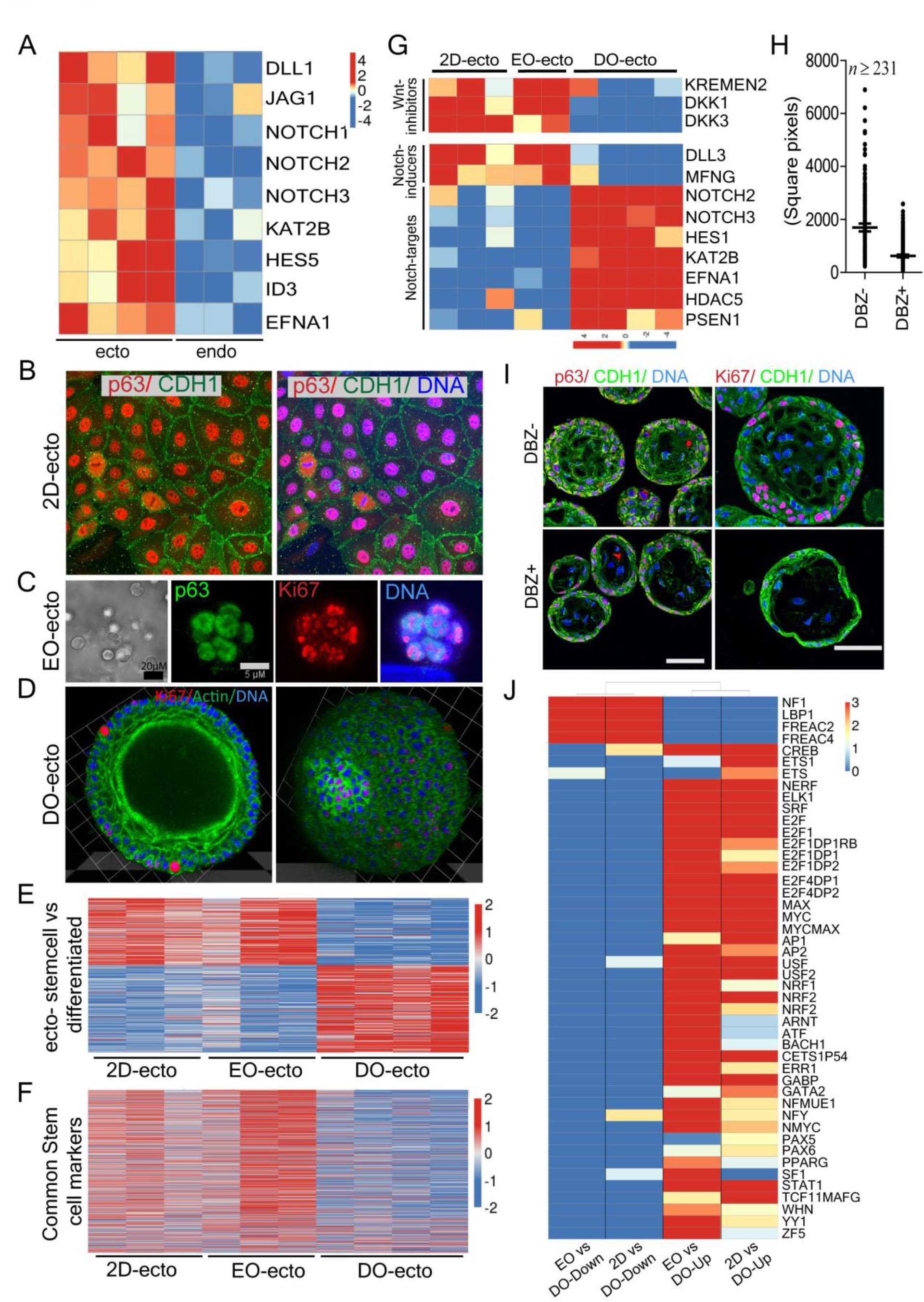
Stemness and differentiation of ectocervix depend on Wnt antagonist, Notch and EGFR signaling. (**A**) Expression analysis of differentially regulated genes in human ecto-vs endocervical organoids. Notch-related genes are expressed at higher levels in ectocervix. Columns = biological replicates. (**B**) Confocal images of 2D human ectocervical stem cell cultures immunolabeled for progenitor cell marker p63 and epithelial cadherin (CDH1). (**C, D**) 3D reconstruction of whole-mount confocal images of 3 day-old early ectocervical organoids labeled for p63 and Ki67 (**C**) and a two-week-old differentiated ectocervical organoid labeled for Ki67 and actin (phalloidin) (**D**). (**E, F**) Heatmaps of differentially regulated genes in 2D as well as early and corresponding differentiated ectocervical organoid cultures (**E**) and genes frequently upregulated in stem cells (**F**) (details see Methods section). (**G**) Heatmap of selected differentially expressed genes showing increased Wnt inhibitors and Notch inducers in 2D cultures and early organoids from ectocervix, in contrast to Notch activation-associated genes in differentiated organoids; columns = biological replicates.(**H**) Quantification of the area of human ectocervical organoids grown in the presence or absence of γ-secretase inhibitor (DBZ), *n* = number of organoids, representative of 3 biological replicates; data represent mean ± s.e.m. (**I**) Confocal images of human ectocervical organoids immunolabeled for CDH1, Ki67 or p63. Inhibition of Notch activation by DBZ prevents differentiation and reduces proliferation. *n* = 3 biological replicates. (**J**) Heatmap showing GSEA enrichment–log10(p-value) of GSEA revealing enrichment of genes upregulated in stem cells regulated by transcription factors downstream of Notch, EGFR-RAS-MAPK target genes signaling among genes upregulated in 2D and EO ectocervical organoids, while the RAS antagonistic NF1 pathway is enriched among genes highly expressed in differentiated ectocervical organoids.

A survey of genes that are upregulated in ectocervical stem cells compared to differentiated cells revealed high expression of the Notch ligands Delta-Like Ligand 3 (DLL3) and Manic Fringe (MFNG), the latter facilitating binding of DLL to the Notch receptor (Fig. 3G). In contrast, differentiated cells expressed higher levels of Notch 2 and Notch 3 receptors as well as their targets, including the transcription factor HES1 and Presenilin 1 (PSEN1), a core component of γ-secretase (Fig. 3G). Ectocervical stem cells also showed highly upregulated expression of the WNT antagonists DKK1, DKK3 and the DKK receptor KREMEN2 (Fig. 3G). Concordantly, inhibition of Notch activation using the γ-secretase inhibitor DBZ reduced organoid growth (Fig. 3H, Fig. S6D), as these organoids failed to differentiate and stratify (Fig. 3I). Thus the ectocervical stem cells act as Notch signal-sending cells, while the differentiated cells show the signature of Notch signal-receiving cells, leading to the trans-activating interaction that facilitates differentiation and ultimately epithelial stratification.

Further, gene set enrichment analysis (GSEA) revealed that genes regulated by several transcription factors downstream of Notch ligand and EGF receptor (EGFR)-RAS-MAPK signalling were highly enriched among genes upregulated in ectocervical stem cells, including AP1 ^13,14^, CREB, ETS, new ETS-related factor (NERF), ELK1, E2F, SRF, MYC and YY1 ^15-18^ (Fig. 3J). The two pathways function together to regulate proliferation and differentiation, with the EGFR pathway promoting the expression of Notch DLL ligands ^19^. On the other hand, genes belonging to the RAS antagonistic NF1 pathway ^20^ were enriched in genes highly expressed in differentiated cells. Together, these observations indicate that the Wnt antagonists together with EGFR and Notch-inducing pathways regulate ectocervical stemness and differentiation.

### The emergence of squamous metaplasia from quiescent KRT5+ stem cells in the endocervix

We next performed *in vitro* and *in vivo* analysis to determine the cellular origin and mechanism of squamous metaplasia. Primary endocervix-derived cells showed a clear enrichment of KRT5+ and p63+ cells if cultured in 2D in Wnt deficient medium. After transfer to organoid culture conditions, these cells produced only organoids of the squamous type, even in the presence of Wnt3a /RSPO1 (Fig. 4A). However, if primary endocervix-derived cells were grown in 2D in a Wnt proficient medium such cultures contained only a few KRT5+ or p63+ cells and gave rise to columnar organoids in the presence of Wnt. Yet, the absence of Wnt favored the growth of squamous organoids, including the characteristic basal and parabasal p63+ cells (Fig. 4A). Importantly though, endocervical organoids derived from single cells remained columnar even when transferred to Wnt deficient medium, thus excluding the possibility that columnar cells transdifferentiate to the squamous lineage (Fig. 4B). In contrast, primary ectocervical cells grown in 2D with either Wnt proficient or deficient medium give rise only to stratified organoids in Wnt deficient medium (Fig. 4C).

**Figure 4.**
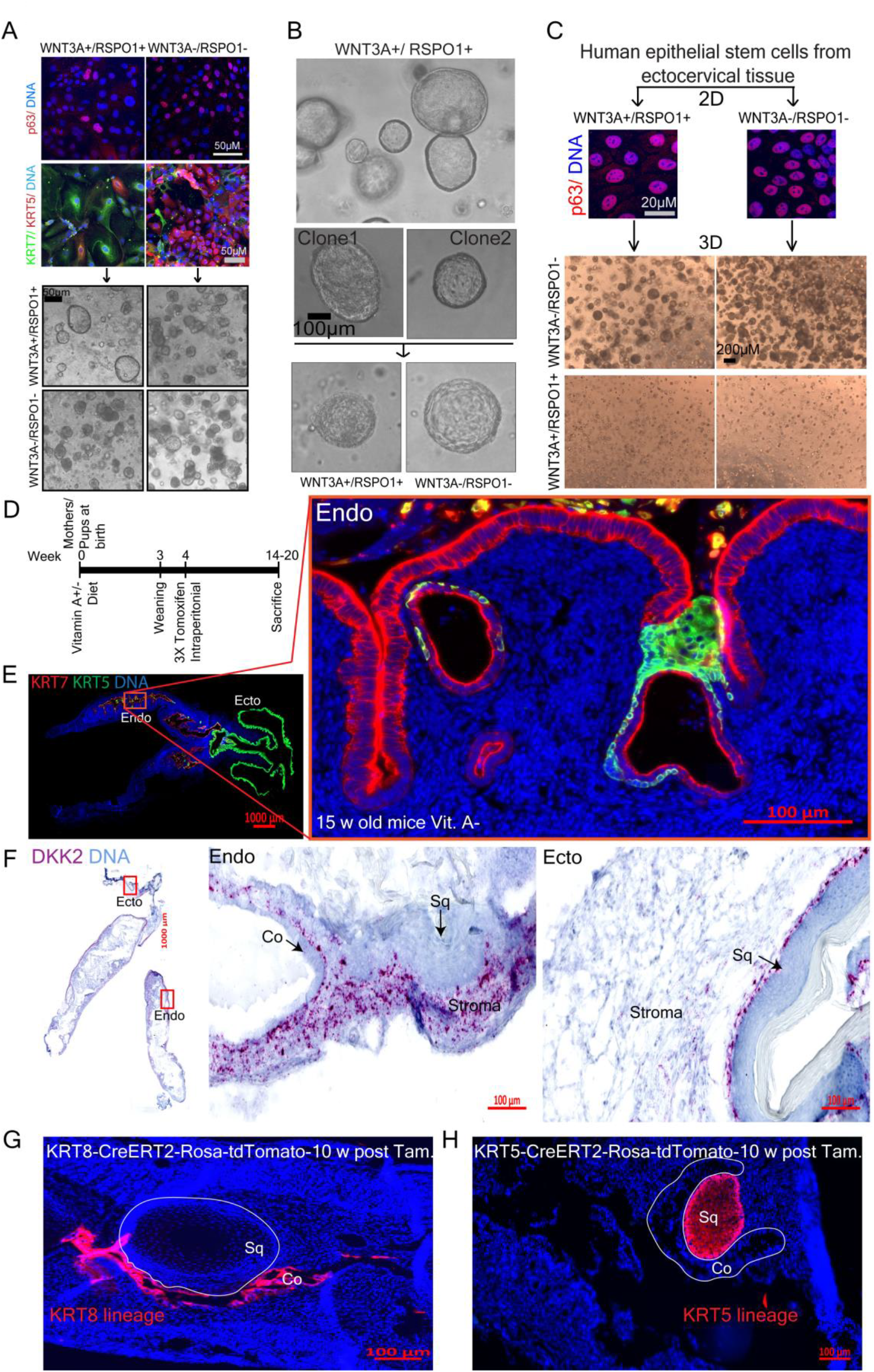
Two distinct stem cells from the endocervix give rise to columnar or squamous stratified lineages depending on the microenvironment. (**A**) Wnt deficient medium enriches for p63^+^/KRT5^+^ endocervical stem cells that only give rise to stratified organoids, while Wnt proficient medium supports both KRT7^+^ and p63^+^/KRT5^+^ cells, which can give rise to columnar or stratified organoids, depending on culture conditions (**B**) Endocervical stem cells that give rise to columnar epithelium are unipotent and fail to transdifferentiate into stratified organoids. Single endocervical organoids were grown in Wnt proficient medium, dissociated into single cells and transferred to Wnt proficient or deficient medium. (**C**) Confocal images of ectocervical epithelial cells grown in 2D. p63^+^ cells are present in Wnt proficient and Wnt deficient media but organoids are formed only in Wnt deficient medium. (**D**) Treatment scheme of Vitamin A deficient diet study in WT and lineage tracing mice. (**E**) Tissue sections from the genital system of C57BL6 mice fed a vitamin A deficient diet for 15 weeks were labeled with antibodies against KRT7 and KRT5. Zoom: outgrowth of subcolumnar KRT5+ stem cells that give rise to squamous metaplastic epithelium in the endocervix. (**F**) Single molecule RNA ISH of tissue from a mouse fed with a vitamin A deficient diet. Expression of DKK2 is enhanced in endocervical stroma. Boxed areas in panel **F** are magnified on right. (**G, H**) Lineage tracing in KRT8CreErt2/Rosa26-tdTomato (**G**) and KRT5CreErt2/Rosa26-tdTomato (**H**) mice fed a vitamin A deficient diet reveals that squamous metaplasia arising in the endocervix is negative for KRT8-tdTomato (**G**) and positive for KRT5-TdTomato (**H**) lineage markers. Fluorescent and brightfield tiled images were acquired with AxioScan imager. Data representative of *n* = 3 biological replicates.

Although the expression of HOX genes, a family of decisive regulators during embryonic development, is largely unknown for the cervix, HOXA11 has previously been associated with cervix development ^21^ and deregulation of HOXB2, HOXB4, and HOXB13 have been implicated in cervical carcinogenesis ^22^. Here we analyzed the pattern of HOX gene expression in the ecto-vs. endocervical organoids (Fig. S7A). Strikingly, we observed substantial differences between the two cultures, supporting the notion that the two tissue types represent different biological lineages in the cervix.

To further consolidate the lineage properties of stratified and columnar epithelial cells and spatial changes in the microenvironment, we performed lineage tracing, single cell RNA-ISH and IHC in a mouse model of squamous metaplasia induced by retinoid depletion ^23^. These retinoid-depleted mice showed an upregulation of DKK2 gene expression in the stroma of the endocervix and uterine horns (here also referred to as endocervix), and the emergence of subcolumnar quiescent KRT5+ cells that eventually developed into metaplastic squamous stratified epithelium (Fig. 4D-F, S7B, compare to Fig. 2F and S4H). However, production of the Wnt target Axin2, which is normally expressed in the endocervix, remained unaltered in these mice, while KRT8 and KRT7 expression were restricted to the columnar epithelium (Fig. 4E, Fig. S7C-E). Further, by performing lineage tracing analysis in retinoid-depleted KRT5CreErt2/Rosa26-tdTomato and KRT8CreErt2/Rosa26-tdTomato mice, we confirmed that KRT8+ cells give rise to columnar epithelium while KRT5+ cells give rise to squamous metaplasia in the endocervix (Fig. 4G-H). Together, these data demonstrate that the endocervix harbors two distinct, unipotent stem cell populations with the potential to develop columnar or stratified lineages, respectively. Which one is activated thus appears to depend on the microenvironment and the opposing Wnt-related signals in particular. While Wnt agonists support the formation of columnar epithelium, the local upregulation of Wnt antagonist in the stroma drives the proliferation of quiescent KRT5+ reserve cells to cause squamous metaplasia.

### Cellular origins of cervical squamous and adenocarcinomas

A number of studies have shown that adult stem cells are susceptible to transformation and often constitute the cells of origin for a variety of cancers ^24^. The origin of ADC and SCC is controversial and uncertain. Here we assessed the expression signatures of squamous and columnar cervical organoids to determine the cells of origin of cervical cancers. We retrieved publically available mRNA expression data for 302 cervical cancers from The Cancer Genome Atlas (TCGA, http://cancergenome.nih.gov/). We used the similarity of differential gene expression profiles between ectocervical squamous and endocervical columnar organoids to those between cancer samples to classify the latter into squamous-like, columnar-like or undetermined cases (Fig.5A, Table S1 and S3, Methods). We found that cancers classified as squamous-like matched a histological diagnosis of SCC in all cases (n=111) while for those classified as columnar-like we found 48/77 matching a histological diagnosis of ADC (Fig. 5B). For 111 cases histologically diagnosed as SCC and 3 ADC, we could not determine a clear classification and 29 SCC were assigned to the columnar-like group by our classifier. Importantly, cancer samples classified as columnar-like were mainly KRT5^low^, KRT7^high^ and p63^low^, while samples in the squamous-like and undetermined group were mainly KRT5^high^ and p63^high^ with mixed KRT7 status (Fig. 5A), suggesting that the undetermined group could consist of SCCs within or outgrown into columnar endocervix, leading to the presence of contaminating endocervical columnar KRT7+ cells in the samples.

**Figure 5.**
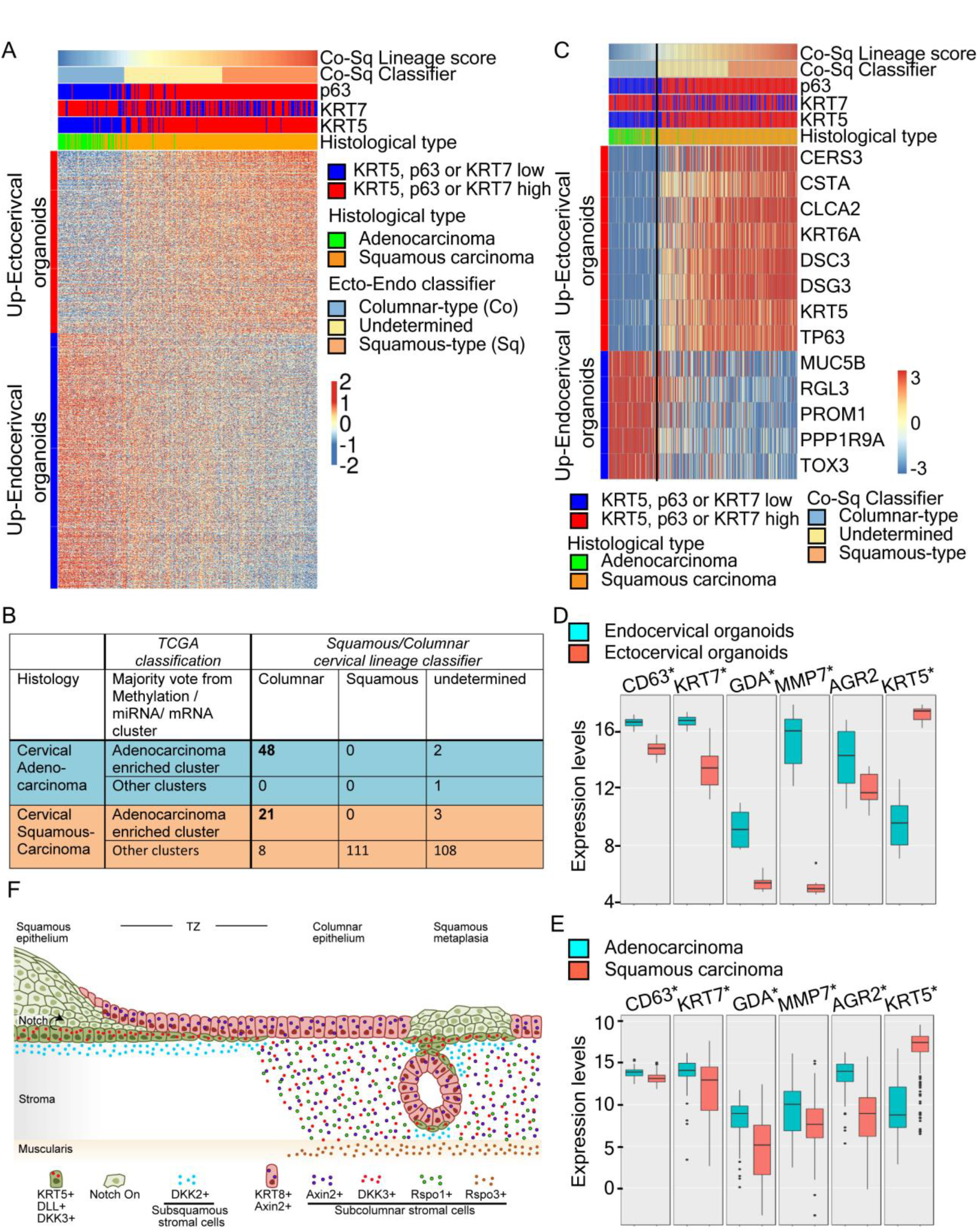
Cervical squamous carcinomas originate from KRT5^+^ and adenocarcinomas from KRT7^+^ stem cells. (**A**) Expression profiles of SCC and ADC correlate well with genes differentially expressed between ecto-and endocervical organoids. (B) Classification of cancer samples based on majority voting from hierarchical mRNA and miRNA or methylation status clustering suggests that 29 samples are histologically incorrectly diagnosed as squamous carcinoma. (**C**) Heatmap showing the mean-substracted expression for selected bimodal genes in cancer samples that are differentially expressed in squamous and columnar organoids. Colour denotes fold-change from mean gene expression across all samples. (**D, E**) Expression profiles of proposed squamocolumnar junction markers together with KRT5 in 302 cervical cancer samples (**D**) and in cervical organoids (**E**). Expression of these markers is higher in endocervical organoids (*n*=6) and ADCs (*n*=51) compared to ectocervical organoids (*n*=10) and SCCs (*n*=251), in contrast to KRT5 expression; * = p<0.05. (**F**) Model depicting the KRT5^+^ and KRT7^+^ stem cell organization and Wnt/Notch microenvironment in TZ and during squamous metaplasia.

To validate these results, we also obtained clustering results based on genome-wide methylation, global mRNA and microRNA expression data for the same cancer samples from TCGA. The TCGA cluster containing most ADCs in each of the clustering analyses from those three levels of cellular regulation showed strong overlap with our columnar-like class. Using the majority vote among mRNA, miRNA and DNA methylation clusters (Fig. 5B) we find that 69/77 samples from the columnar-like group are also in the TCGA clusters enriched for ADC, while all other TCGA clusters together contain mainly squamous-like and undetermined samples (228/231) (Fig. 5B and S8). Interestingly, 21/29 cancers defined as columnar-like based on our classifier, but histologically classified as SCC, showed strong similarity to ADC according to TCGA molecular profiles and might therefore be misdiagnosed.

A recent study suggested that only a small population of cells located in the TZ (the so-called squamo-columnar junction (SCJ) cells) express KRT7 and that these are the precursors of both SCC and ADC ^4^. We further investigated the mRNA expression levels in organoids and 302 cervical cancer samples from TCGA with regard to SCJ markers proposed in that study ^4^, as well as KRT5. In contrast to KRT5, expression of the proposed SCJ markers is significantly higher in healthy endocervical organoids as compared to healthy ectocervical organoids and the same trend is seen in ADC vs. SCC (Fig. 5D-E). This indicates that the reported SCJ cells are not distinct from the endocervical columnar lineage and are not the cells of origin for SCC.

Our study also revealed a set of genes that are differentially expressed between squamous and columnar organoids and show a strong correlation with columnar-like and squamous-like cancers, including MUC5B, KRT5, CSTA, while the proposed SCJ markers KRT7, AGR2 and GDA specifically labeled ADC but not SCC sections (Figure 5C and S9). Thus, the majority of cervical cancers can be divided into two subgroups based on molecular signatures that correlate with signatures of KRT5+ stem cells for squamous or KRT7+/KRT8+ stem cells for columnar cervical epithelia. Together, these results indicate that the cervix harbors two distinct stem cell lineages, reflecting the cells of origin for SCC and ADC, respectively.

## Discussion

The TZs of the mucosal epithelium constitute critical zones of enhanced disposition to infections and carcinogenesis ^25-29^. Revealing the principles of cellular regulation and homeostasis of these tissue regions is key to understanding the impact of intrinsic and extrinsic disturbances, as well as for prospective and therapeutic disease prevention.

We show that distinct microenvironmental conditions and molecular signals from the epithelial and stromal tissues drive the dominance of specific epithelial lineages of the TZ. We reveal Wnt signaling as a key determinant in regulating the homeostasis at borders between two epithelial types. Wnt signaling has been shown to be indispensable for the maintenance and homeostasis of adult stem cells in several mammalian tissues ^11,30^. However, here we show that in the cervix, Wnt signaling stimulated by the underlying stroma drives the columnar lineage while imposing quiescence of squamous lineage-specific stem cells that exist in the same milieu. With the transition to a Wnt repressive microenvironment, these quiescent squamous lineage stem cells are activated and replace the columnar epithelia at the TZ or as an island of metaplasia within the endocervix. Further, the fact that ADC or SCC arise from two distinct stem cell lineages rather than a common cellular origin has important clinical implications for choice of therapy and suggests that preventive ablation of the SCJ alone may not fully eliminate potential cervical cancer precursor cells ^7,8^.

Our data constitute a major conceptual progress in our understanding of how epithelial junctions are maintained in our body. Accordingly, homeostasis at these sites is not maintained by the transition from one epithelial type to another but rather that the adult tissue is composed of different stem cell populations that are retrieved upon extrinsic signals to generate respective cell lineages, forming the adult tissue. This novel concept of homeostasis of the mucosal TZs fits well with other recent observations on the mucosal stem cell identity and may stimulate future investigations with therapeutic relevance.

## Author contributions

CC, RKG and TFM conceived the study; CC, RKG designed and CC, RKG and SK performed experiments;HB conceived and performed the in silico analyses; VB, UK, and HM contributed imaging, mouse breeding, microarray studies, respectively; MM and HH provided human samples; CC and RKG analyzed the data, and wrote the manuscript; HB and TFM revised the manuscript; TFM supervised the study.

## Acknowledgements

The authors would like to thank Marina Drabkina and Christiane Dimmler for excellent technical assistance, Kirstin Hofmann for excellent help with mouse experiments, Ina Wagner for the microarrays, Gesa Rausch and Michael Meyer for assistance with the animal application, Diane Schad for assistance with preparing the graphics and Rike Zietlow for editing the manuscript.

This work was supported by the BMBF through the Infect-ERA project CINOCA (FK031A409A to TFM). The funders had no influence on the study design or analysis of the data.

## Data availability

Microarray data have been deposited in the Gene Expression Omnibus (GEO; www.ncbi.nlm.nih.gov/geo/) of the National Center for Biotechnology Information and can be accessed with the GEO accession number GSE87076.

## Materials and Methods

### Antibodies and Chemicals

The following antibodies and chemicals were used: mouse-anti-p63 (Abcam, # ab375), rabbit-anti-p63 (Abcam, # ab53039), mouse-anti-E-Cadherin (BD Biosciences, # 610181), rabbit-anti-Ki67 (Abcam, # ab16667), mouse/rat-anti-Ki67-FITC (eBioscience, # 11-5698), mouse-anti-KRT5 (Sigma, # C-7785), rabbit-anti-KRT5 (Abcam, # ab52635), rabbit-anti-cytokeratin 5-Alexa488 (Abcam, # ab193894), mouse-anti-KRT7 (Santa Cruz, # sc-23876), rabbit-anti-cytokeratin 7 (Abcam, # ab181598), rabbit-anti-cytokeratin 7-Alexa555 (Abcam, # ab209601), rabbit-anti-CSTA (Cystatin A) (Sigma # HPA001031), rabbit-anti-AGR2 (Proteintech, # 12275-1-AP), mouse-anti-MUC5B (Abcam, # ab77995), rabbit-anti-GDA (Sigma # HPA019352), Hoechst (Sigma, #B2261), Draq5 (Cell Signaling, #4085), γ-secretase inhibitor XX (DBZ) (Calbiochem # 565789) and p38 inhibitor SB202190 (Sigma, # S7067). Secondary antibodies labeled with the fluorochromes Cy2, Cy3 or Cy5 were obtained from Jackson ImmunoResearch Laboratories.

### Mouse experiments

All procedures involving animals were approved by the national legal as well as institutional and local authorities at the Max Planck Institute for Infection Biology. Wild-type C56BL6, KRT5CreErt2 ^31^ and KRT8CreErt2 ^32^ mice were obtained from the Jackson Laboratory. These strains were bred to Rosa-tdTomato ^33^ mice in order to generate mice expressing a fluorophore in Cre-expressing cells. For lineage analysis for the cell of origin of Krt5+ or KRT8+ cells, Cre recombinase was induced in female mice by administering tamoxifen (Sigma) intraperitoneally at 0.25 mg g^−1^ body weight in 50 μl corn oil at week 4 on three consecutive days. Mice were euthanized at 14-20 weeks and the genital tracts removed for further analysis.

### Depletion of retinoid signaling in mice using Vitamin A-deficient diet

At birth experimental mice and their mothers were placed on a vitamin A-deficient test diet (SAFE, U8978P-0074) or control diet with added Vitamin A at physiological levels of 6 IU/g (SAFE, U8978P-0075) following a protocol developed for BALB/c mice ^23^. Littermates were weaned at 3 weeks of age and maintained on the deficient or control diet for a period of 14-20 weeks before being euthanized for further analysis.

### Mouse cervical medium

Cervical cells were cultured in ADF medium (Invitrogen, # 12634) supplemented with 12 mM HEPES (Invitrogen, # 15630-056), 1% GlutaMax (Invitrogen, # 35050-038), 1% B27 (Invitrogen, # 17504-044), 1% N2 (Invitrogen, # 17502048), 50 ng/ml mouse epidermal growth factor (EGF) (Invitrogen, # PMG8043), 100 ng/ml mouse noggin (Peprotech, # 250-38-100), 100 ng/ml human fibroblast growth factor (FGF)-10 (Peprotech, # 100-26-25), 1.25 mM N-acetyl-L-cysteine (Sigma, # A9165-5G), 10 mM nicotinamide, (Sigma, # N0636), 2 μM TGF-β R Kinase Inhibitor IV (Calbiochem, # 616454), 10 μM ROCK inhibitor (Y-27632) (Sigma, # Y0503), 1% penicillin/streptomycin (Gibco, # 15140-122) with or without 25% Wnt3A-and 25% R-spondin1-conditioned medium, as described in Willert et al ^34^ and Farin et al ^35^.

### Human ectocervical (Wnt deficient) medium

Consisted of ADF, 12 mM HEPES and 1% GlutaMax, supplemented with 1% B27, 1% N2, 0.5 μg/ml hydrocortisone (Sigma, # H0888-1G), 10 ng/ml human EGF (Invitrogen, # PHG0311), 100 ng/ml human noggin (Peprotech; # 120-10C), 100 ng/ml human FGF-10 (Peprotech, # 100-26-25), 1.25 mM N-acetyl-L-cysteine, 10 mM nicotinamide, 2 μM TGF-β R kinase Inhibitor IV, 10 μM ROCK inhibitor (Y-27632), 10 μM forskolin (Sigma, F6886) and 1% penicillin/streptomycin.

### Human endocervical (Wnt proficient) medium

Consisted of ADF, 12 mM HEPES, 1% GlutaMax, supplemented with 1% B27, 1% N2, 10 ng/ml human EGF, 100 ng/ml human noggin, 100 ng/ml human FGF-10, 1.25 mM N-acetyl-L-cystein, 10 mM nicotinamide, 2 mM TGF-β R kinase Inhibitor IV and 10 μM ROCK inhibitor (Y-27632) with 25% Wnt3A-and 25% R-spondin1 conditioned medium.

### Epithelial stem cell isolation from human and mouse cervix

Human ecto-and endocervix samples were provided by the Department of Gynecology, Charité University Hospital, Berlin, Germany. Scientific usage of the samples was approved by the ethics committee of the Charité University Hospital, Berlin (EA1/059/15); informed consent to use their tissue for scientific research was obtained from all subjects. Only anatomically normal tissues were used, within 2-3 h after of removal. Mouse cervix was removed from euthanized 4-8 week old healthy female wild type BALB/c mice (from Charles River) immediately preceding the isolation of the cells. Tissue samples were washed thoroughly in sterile PBS (Gibco, # 14190-094) and minced with surgical scissors. Minced tissue was incubated in 0.5 mg/ml collagenase type II (Calbiochem, # 234155) for 2.5 h at 37 °C in a shaker incubator. Tissue and dissociated cells were pelleted by centrifugation (5 min at 1000 g, 4 °C), supernatant discarded, cells resuspended in TrypLE express (Gibco, # 12604021) and incubated for 15 min at 37 °C in a shaker incubator. After dissociation, the cell and tissue pellet was resuspended in ADF (Invitrogen) medium and passed through a 40 μM cell strainer (BD Falc, # 352340) to separate the single dissociated cells from tissue pieces. Cells were pelleted by centrifugation (5 min at 1000xg, 40 °C), resuspended in either human ecto-or endocervical or mouse cervical medium and cultured either directly as organoids or in 2D.

### Human epithelial stem cell culture and maintenance in 2D

Human epithelial stem cells isolated from the tissue were resuspended in either ecto-or endocervical medium and plated in collagen-coated tissue culture flasks. Cells were incubated at 37 °C, 5% CO2 in a humidified incubator. Once they reached 70-80% confluence, cells were detached using TrypLE Express, and centrifuged at 1000xg for 5 mins at 40 °C. The cells were then used for culturing organoids or for maintenance of 2D stem cells. 2D stem cells were maintained by seeding the 2D cells from P1 into tissue culture flasks containing lethally irradiated J2-3T3 fibroblast feeder cells in ecto-or endocervical medium. Medium was replaced and irradiated fibroblasts added every 4 days until the colonies reached a confluence of 60-70%, at which stage they were detached and reseeded onto freshly irradiated feeders at a 1:5 ratio or cryopreserved for later use.

### Organoid culture and maintenance

Cells isolated from tissue or the stem cells grown in 2D culture were mixed with 50 μl of ice-cold Matrigel (BD, # 356231) at a density of 20,000 cells, and the Matrigel droplet was placed in a pre-warmed 24-well plate and allowed to polymerize for 10 min at 37 °C. The Matrigel droplet was then overlaid with 500 μl of pre-warmed human ecto-or endocervical medium. Cultures were kept at 37 °C, 5% CO2 in a humidified incubator for 2-3 weeks and medium replaced every four days. For passaging the organoids, Matrigel was first dissolved by adding 1 ml of ice-cold ADF and pipetting up and down 5 times. Organoids were collected in a 15 ml Falcon tube and a further 4 ml of ice-cold ADF medium was added and organoids resuspended well to completely dissolve the Matrigel, followed by centrifugation at 300xg for 5 mins at 4°C. Medium was discarded and the ectocervical and mouse organoids were incubated with 1 ml of TrypLE Express for 30 min at 37 °C followed by mechanical fragmentation using a fire-polished glass Pasteur pipette by vigorous pipetting (8–10 times) to generate single cells. The single cells were then seeded at a 1:10 ratio back into Matrigel for expanding and culturing. For the endocervical organoids, after centrifugation organoids were subjected to mechanical fragmentation as described above to generate fragments that were seeded back into Matrigel at 1:5 ratio. Matrigel was allowed to polymerize for 10 min at 37°C, overlaid with pre-warmed medium and cultured as described above.

### Organoid forming ability

Stem cells were counted and a defined number resuspended in 50 μl Matrigel to generate organoids as described above. Between 2-3 weeks after plating images were taken of the whole well and the number and area of organoids formed were determined using ImageJ to calculate organoid forming efficiency.

### Immunofluorescent histochemistry

Organoids were washed five times with cold PBS to remove Matrigel before fixing with 4% paraformaldehyde for 1 h at room temperature (RT) followed by washing with PBS twice. Organoids were then subjected to dehydration in an ascending ethanol series followed by isopropanol and acetone for 20 min each. The dehydrated organoids were paraffin-embedded and 5 μM sections cut on a Microm HM 315 microtome. Mouse and human tissues were extensively washed with PBS and fixed using 4% PFA overnight at RT. Samples were subjected to dehydration in an ascending ethanol series followed by isopropanol and xylene (60 min each) followed by paraffinization using a Leica TP1020 tissue processor. The tissue was embedded and 5 μM sections cut on a microtome. For immunostaining, paraffin sections were deparaffinized and rehydrated, followed by treatment with antigen retrieval solution (Dako, # S1699). Sections were blocked using blocking buffer (1% BSA and 2% FCS in PBS) for 1 h at RT. Primary antibodies were diluted in blocking buffer and incubated for 90 mins at RT followed by five PSB washes before 1 h incubation with secondary antibodies diluted in blocking buffer along with Hoechst or Draq5. Sections were washed with PBS five times and mounted using Mowiol. Images were acquired with a Leica TCS SP8 confocal microscope.

Fresh epithelial isolates were grown on collagen-coated coverslips in 2D and fixed with 4% paraformaldehyde for 30 min at RT. Cells were permeabilized and blocked with 0.5% Triton X-100 and 1% BSA in PBS. Primary antibodies were diluted in 1% BSA in PBS and incubated for 1 h at RT followed by three washes in PSB-T (0.1% Tween 20 in PBS), followed by 1 h incubation with secondary antibodies diluted in 1% BSA in PBS along with Hoechst or Draq5. Coverslips were washed three times with PBS-T and once with PBS and mounted using Mowiol. Images were acquired on a Leica TCS SP8 confocal microscope. Images were processed with Adobe Photoshop; 3D reconstruction was done with the Volocity 6.3 software package (Perkin Elmer).

### Whole mount staining

Matrigel was removed from the organoids by extensive washing with ice-cold PBS prior to fixation (4x 45 min) and allowed to settle by gravity to maintain the 3D structure. Organoids were then fixed using pre-warmed (37 °C) 3.7% PFA for 1 h at RT followed by three PBST washes. Permeabilization and blocking was performed overnight at 40 °C using 5% donkey serum, 1% FCS, 0.05% Tween20, 2% Triton X-100, 0.02% sodium azide in PBS. Organoids were incubated with primary antibodies diluted in blocking buffer (5% donkey serum, 1% FCS, 0.25% Triton X-100, 0.02% sodium azide in PBS) at 4°C for 3-5 days followed by three PBST washes for 45 min each at RT. Next, organoids were incubated with secondary antibodies diluted in blocking buffer for two days at 4°C followed by one PBST wash for 45 min and three washes with PBS containing 5% glycerol for 45 min each. Organoids were then carefully transferred to an ibidi μ-slide (# 81822) together with some PBS and glycerol solution and Z stack images were acquired with a confocal microscope and image processing and 3D reconstructions were done using Volocity 6.3 software.

### Single-molecule RNA in situ hybridization (RNA-ISH)

For single molecule RNA in situ labelling, paraffin embedded 10 μM tissue sections were used with RNAscope 2.5 HD Red Reagent kit (Advanced Cell Diagnostics). Hybridizations were performed according to the manufacturer’s protocol. In each experiment, positive (PPIB) and negative (DapB) control probes were used as per the manufacturer’s guidelines. Tiled bright field images were obtained with Axio Scan.Z1 tissue imager (Zeiss). Images were further processed with Zen 2.3 (Blue edition) image analysis software and further compiled using Adobe illustrator.

### RNA isolation and quality control

Microarrays were hybridized for human ectocervical cells cultured in 2D in Wnt-deficient medium (n=3 biological replicates from 2 human donors) or as organoids (EO: n=3 biological replicates from 3 human donors, DO: n=4 biological replicates for 4 human donors), human endocervical cells cultured in 2D Wnt-proficient medium (n=3 biological replicates from 3 human donors) or as DO organoids (n=3 biological replicates from 3 human donors), as well as mouse cervical EO and DO organoids cultured in Wnt-proficient or-deficient medium, respectively (n=2 biological replicates per condition). In the absence of any pre-existing knowledge on expected effect sizes sample sizes were selected based on available samples. Cells and organoids were pelleted and resuspended in 1 ml Trizol (Life Technologies) and RNA was isolated according to the manufacturer’s protocol. Quantity of RNA was measured using a NanoDrop 1000 UV-Vis spectrophotometer (Kisker) and quality was assessed by Agilent 2100 Bioanalyzer with an RNA Nano 6000 microfluidics kit (Agilent Technologies).

### Microarray expression profiling and Data analysis

Microarray experiments were performed as single-color hybridizations on custom whole genome human 8×60k Agilent arrays (Design ID 048908) and Agilent Feature Extraction software was used to obtain probe intensities. The extracted single-color raw data files were background corrected, quantile normalized and further analyzed for differential gene expression using R ^36^ and the associated BioConductor package LIMMA ^37^ (Table S2). Microarray gene expression comparisons between groups were performed using unpaired tests for all human comparisons. R was also used for all statistical analyses unless stated otherwise. Mann-Whitney-U test was used for comparisons of gene expression in SCJ marker genes with a threshold of p<0.05. Microarray data have been deposited in the Gene Expression Omnibus (GEO; www.ncbi.nlm.nih.gov/geo/) of the National Center for Biotechnology Information and can be accessed with the GEO accession number GSE87076.

The signature of differentially expressed genes between ectocervical 2D/EO vs DO organoids was selected from all genes with a false discovery rate (FDR) < 0.05 and log2 fold change < −1.5 or > 1.5 in any of the two comparisons (2D vs. DO or EO vs. DO) and the largest absolute fold change from both comparisons and possible replicate probes was taken for each gene.

### Analysis of stem cell related genes

Raw data from different microarray data sets obtained from adult tissue stem cells (SC) cultured on feeder cells and corresponding differentiated cells from air-liquid interface (ALI), Matrigel or self-assembly sphere (SAS) were downloaded from GEO (GSE57584, GSE66115, GSE69453, GSE65013, GSE32606, GSE69429, GSE49292) and normalized together using method ‘RMA-sketch’ with Affymetrix Power Tools. We assessed differentially expressed genes between SC and corresponding differentiated cell cultures for normal esophagus, Barrett’s esophagus, gastric cardia, duodenum, jejunum, ileum, colon ascendens, colon transversum, colon descendens, KRT5+ and KRT7+ fetal esophageal cells, fallopian tube, nasal turbinated epithelium, tracheobronchial epithelium and distal airway epithelium. We selected stem cell-related genes as those genes with significant (adjusted p-value < 0.05) up-or down-regulation (abs(logFC) > 1) in at least 5 out of 18 comparisons (Table S1).

### Gene Set Enrichment Analysis (GSEA)

We performed a pre-ranked GSEA analysis using GSEA software v2.1.0 ^38,39^ obtained from http://software.broadinstitute.org/gsea. The t-statistics from comparisons of ectocervical organoids (2D vs. Differentiated organoids or Early organoids vs. Differentiated organoids) were used to rank probes and enrichment of MSigDB Motif gene sets [http://software.broadinstitute.org/gsea/msigdb] (c3.all.v5.1.symbols.gmt) was computed using standard settings, collapsing probe sets within genes using the Max_probe method and using 1000 permutations. For further analysis we kept only motif gene sets that were significant in at least one of the up or down regulated genes in the two comparisons mentioned above at FDR < 5%. For the heatmap visualization, we chose the smallest p-value for motif gene sets referring to the same transcription factor use the negative log10 of this value for visualization.

### Cervical cancer data

Expression data (Level 3 processed RNASeq_v2) was obtained for 302 unique samples with available histological diagnosis from The Cancer Genome Atlas (TCGA) data portal (https://gdc-portal.nci.nih.gov/). This data was generated within the Cervical Squamous Cell Carcinoma and Endocervical Adenocarcinoma project (TCGA-CESC) and is a superset of the published cohort 40. Per gene expression levels were extracted from "*.rsem.genes.normalized_results" files using custom scripts. Public clinical sample annotations for t0hose samples were also obtained from the same source. Aggregated features including clustering results based on DNA methylation, mRNA and microRNA expression was obtained from the Cervical and Endocervical Cancer (CESC) project Firehose site of TCGA 41. For the details on the majority vote see Fig. S8. To classify samples into squamous-like and columnar-like classes, the gene expression levels were log2 transformed and Z-score was applied to make genes comparable. A squamous vs columnar organoid signature was defined based on the fold changes between ectocervical squamous and endocervical columnar differentiated organoids for 2,834 genes with FDR < 0.05 and absolute log2 fold change > 1, selecting the probe with the lowest p-value for each gene. Spearman correlation coefficients (referred to as Co-Sq Score) were computed between Z-scored gene expression values from each cancer sample and the corresponding fold change for the same gene from the squamous vs columnar organoid signature. We defined samples with Co-Sq Score > 0.2 as squamous-like, those with <-0.2 as columnar like and all other as ‘undetermined’ (Fig S8).

Applying the same procedure to 1,000 random sets of genes of the same size with the same fold changes produced sample correlation coefficients generally lower than |0.06|. Thresholds for classification of samples into KRT5-high/low and KRT7-high/low as well as TP63 high/low classes were selected manually to separate the highest cluster from all other samples (Fig. S8). For simplicity, we combined all diagnoses with an adenoma component (Endocervical Adenocarcinoma, Endometrioid Adenocarcinoma, Mucinous Adenocarcinoma and Adenosquamous Carcinoma) into Cervical Adenocarcinoma (Tables S1 and S3).

### Code Availability

All R code used for generating analyses used in this publication is available from the authors on request.

